# A comprehensive single cell data analysis of in lymphoblastoid cells reveals the role of Super-enhancers in maintaining EBV latency

**DOI:** 10.1101/2022.08.10.503552

**Authors:** Bingyu Yan, Chong Wang, Srishti Chakravorty, Zonghao Zhang, Simran D. Kadadi, Yuxin Zhuang, Isabella Sirit, Yonghua Hu, Minwoo Jung, Subhransu Sahoo, Luopin Wang, Kunming Shao, Nicole L. Anderson, Jorge L. Trujillo-Ochoa, Xing Liu, Matthew R. Olson, Behdad Afzali, Bo Zhao, Majid Kazemian

**Author notes:** These authors contributed equally (Bingyu Yan, Chong Wang, Srishti Chakravorty, Zonghao Zhang, Simran D. Kadadi).

## Abstract

We probed the lifecycle of EBV on a cell-by-cell basis using single cell RNA sequencing (scRNA-seq) data from nine publicly available lymphoblastoid cell lines (LCL). While the majority of LCLs comprised cells containing EBV in the latent phase, two other clusters of cells were clearly evident and were distinguished by distinct expression of host and viral genes. Notably, both were high expressors of EBV *LMP1*/*BNLF2* and *BZLF1* compared to another cluster that expressed neither gene. The two novel clusters differed from each other in their expression of EBV lytic genes, including glycoprotein gene *GP350*. The first cluster, comprising *GP350*^−^*LMP1*^*hi*^ cells, expressed high levels of *HIF1A* and was transcriptionally regulated by HIF1-α. Treatment of LCLs with Pevonedistat, a drug that enhances HIF1-α signaling, markedly induced this cluster. The second cluster, containing *GP350*^+^*LMP1*^*hi*^ cells, expressed EBV lytic genes. Host genes that are controlled by super-enhancers (SEs), such as transcription factors *MYC* and *IRF4*, had the lowest expression in this cluster. Functionally, the expression of genes regulated by MYC and IRF4 in *GP350*^+^*LMP1*^*hi*^ cells were lower compared to other cells. Indeed, induction of EBV lytic reactivation in EBV^+^ AKATA reduced the expression of these SE-regulated genes. Furthermore, CRISPR-mediated perturbation of the *MYC* or *IRF4* SEs in LCLs induced the lytic EBV gene expression, suggesting that host SEs and/or SE target genes are required for maintenance of EBV latency. Collectively, our study revealed EBV associated heterogeneity among LCLs that may have functional consequence on host and viral biology.

**Importance:** Epstein-Barr virus (EBV) establishes a life-long latency program within host cells. As such, EBV immortalized lymphoblastoid cells (LCLs) often carry the latent EBV genome and only a small percentage of LCLs containing lytic EBV. However, the cellular programs that distinguish latent from lytic cells and the heterogeneity of cells in latent or lytic phases remains poorly explored. To explore these unknowns, we reanalyzed publicly available single cell RNA-seq data from nine LCLs. This approach permitted the simultaneous study of cells in both latent and lytic phases. We identified three cell populations with distinct lytic/latent activity and further characterized the transcriptomes of these cells. We also identified a new role of super-enhancers in regulating EBV lytic replication. Collectively, our studies revealed EBV associated heterogeneity among LCLs that contribute to EBV life cycle and biology.

## Introduction

Epstein-Barr virus (EBV) is the first oncogenic human DNA virus discovered more than 50 years ago (1). EBV causes ∼200,000 cases of diverse cancers every year (2), including lymphomas, nasopharygeal carcinoma and gastric adenocarcinomas (3, 4). Most EBV infections occur early in life and are transmitted through saliva. EBV first infects oral epithelial cells and then B lymphocytes in the oral epithelium. EBV persists in memory B cells for life in a latent phase, so these cells express minimum EBV genes under host immune surveillance. However, when host immunity is impaired, for example by immunosuppressive treatment or HIV infection, EBV in infected B cells can enter type III latency where six EBV nuclear antigens (EBNAs), three latent membrane proteins, and a few noncoding RNAs and microRNAs, are expressed (3, 5). This can result in lymphoproliferative diseases or lymphomatous transformation (6). EBV in infected memory B cells can also enter a lytic phase to actively produce live virus. During EBV lytic replication, immediate early genes RTA and ZTA encoded by *BRLF1* and *BZLF1* genes, respectively, are first expressed. These are transcription factors (TFs) that turn on the expression of genes necessary for viral DNA replication and structural proteins, including viral membrane protein gp350, that binds to human B cell EBV receptor CD21, to assemble virions (7).

*In vitro*, EBV infection of primary B lymphocytes leads to the establishment of lymphoblastoid cell lines (LCLs) (8). LCLs express EBV type III latency genes, the same genes seen in some EBV malignancies, including post-transplant lymphoproliferative disease and AIDS CNS lymphomas. Therefore, LCLs are an important model system to study EBV oncogenesis. Genetic studies have found that EBNA1, 2, LP, 3A, 3C and LMP1 are essential for EBV-mediated growth transformation (9, 10). EBNA1 tethers EBV episomes to host DNA (11-13). EBNA2 and LP are the major EBV transcription activators that activate expression of key oncogenes, including MYC (14, 15). EBNA3A and 3C repress expression of p16^INK4A^ and p14^ARF^, to overcome senescence and BIM to avoid apoptosis (16-18). LMP1 activates NF-κB to provide survival signals (19).

EBV infection significantly alters chromatin topology and function at EBV-interacting genomic loci of host cells (20). This alteration could be mediated by EBV-encoded transcription factors (21, 22) or via interaction between EBV episomes and the host genome (23), and may also depend on the EBV latency program (24, 25). Super-enhancers (SEs) are critical regions of mammalian genomes comprised of clusters of enhancers bound by arrays of transcription factors (26). Viral transcription factors and host NF-κB subunits can form EBV SEs (22) with markedly high and broad histone 3 lysine 27 acetylation (H3K27ac) (27). These SEs are linked to many genes essential for LCL growth and survival, including MYC and IRF4, and their perturbation pauses LCL growth and causes cell death (28, 29). We have previously shown that EBV episomes physically interact with SE-containing genomic host loci in EBV-transformed lymphoblastoid cells (23). However, the consequences of these interactions and the effects of perturbations at these SE-containing loci on the EBV life cycle remains unexplored.

High throughput sequencing technologies can aid to dissect mechanisms underlying host-virus interactions (4, 23, 30, 31). However, the heterogenous nature of virally infected cells is an impediment to precisely probing the phases of virus life cycle and their effect on host genes in individual cells. Recent advances in single-cell transcriptomics have enabled successful resolution of tissue/cell heterogeneity in several species. Since these technologies agnostically capture both host and infecting viral sequences, they have also been utilized to explore host-virus interactions at a single cell level (32-34). Leveraging this feature here, we sought to identify the determinants of EBV latency in lymphoblastoid cells. Using single cell transcriptomics, we identified distinct clusters marked by differences in expression of *GP350* and *LMP1*. Cells expressing high levels of *LMP1*, but not *GP350*, demonstrated high HIF1-α activity and could be induced by a HIF1-α stabilizer. Cells co-expressing *GP350* and *LMP1/BNLF2* had significantly reduced expression of SE-containing genes compared to cells containing EBV that was clearly in the latent phase (i.e., *GP350*^−^ cells). Using proof-of-principle SE inactivation experiments, we found that host SEs are necessary for the maintenance of EBV latency. Collectively, our data not only highlight the heterogeneity among LCLs but also identifies common functional themes of the cells and their role in EBV associated biology.

## Results

### Single cell RNA-sequencing analyses resolve LCLs into three distinct populations

To better understand how EBV in infected cells spontaneously enter the lytic life cycle, we analyzed publicly available single cell RNA-sequencing (scRNA-seq) data from nine LCLs from several independent sources (see Methods) (33, 35-39). Briefly, we performed an unbiased integrative analysis across all these LCLs after regressing for potential batch effects, doublets and/or artifacts and known sources of heterogeneity, such as the stage of cell cycle using the Seurat platform (40) (**Figs. S1a-c-** see methods). Unsupervised clustering of all 46,205 cells according to expressions of both host and viral genes at three different resolutions yielded several clusters (**Fig. S1d**). Further examination of these clusters based on the expression of salient EBV genes, including *GP350* and *LMP1*/*BNLF2*, and separation in UMAP space revealed that these clusters fall into three major groups according to the status of EBV gene expression, namely latent, early lytic and full lytic EBV cell clusters. The multiple clusters corresponding to EBV in the latent state were recently examined thoroughly (33). Since our focus was mainly on understanding the biology of EBV lytic life cycle, we combined all the latent cells into one cluster, resulting in three major clusters (**Fig. 1a**). These were evident in all LCL datasets examined (**Figs. S1a-b**). These clusters contained *GP350*^−^*LMP1*^*lo*^, *GP350*^−^*LMP1*^*hi*^ and *GP350*^+^*LMP1*^*hi*^ cells, representing cells with EBV in the latent, early lytic and fully lytic states (**Figs. 1a, S1e**). Approximately 50-100 host genes were differentially expressed in each cluster compared to all other clusters (**Figs. 1b, S1f** and **Table S1**). Consistently, *GP350*^+^*LMP1*^*hi*^ cells was the cluster expressing the most EBV genes, while *GP350*^−^*LMP1*^*lo*^ cells represented the cluster showing the lowest expression of EBV genes (**Figs. 1b-d**).

**Figure 1.**
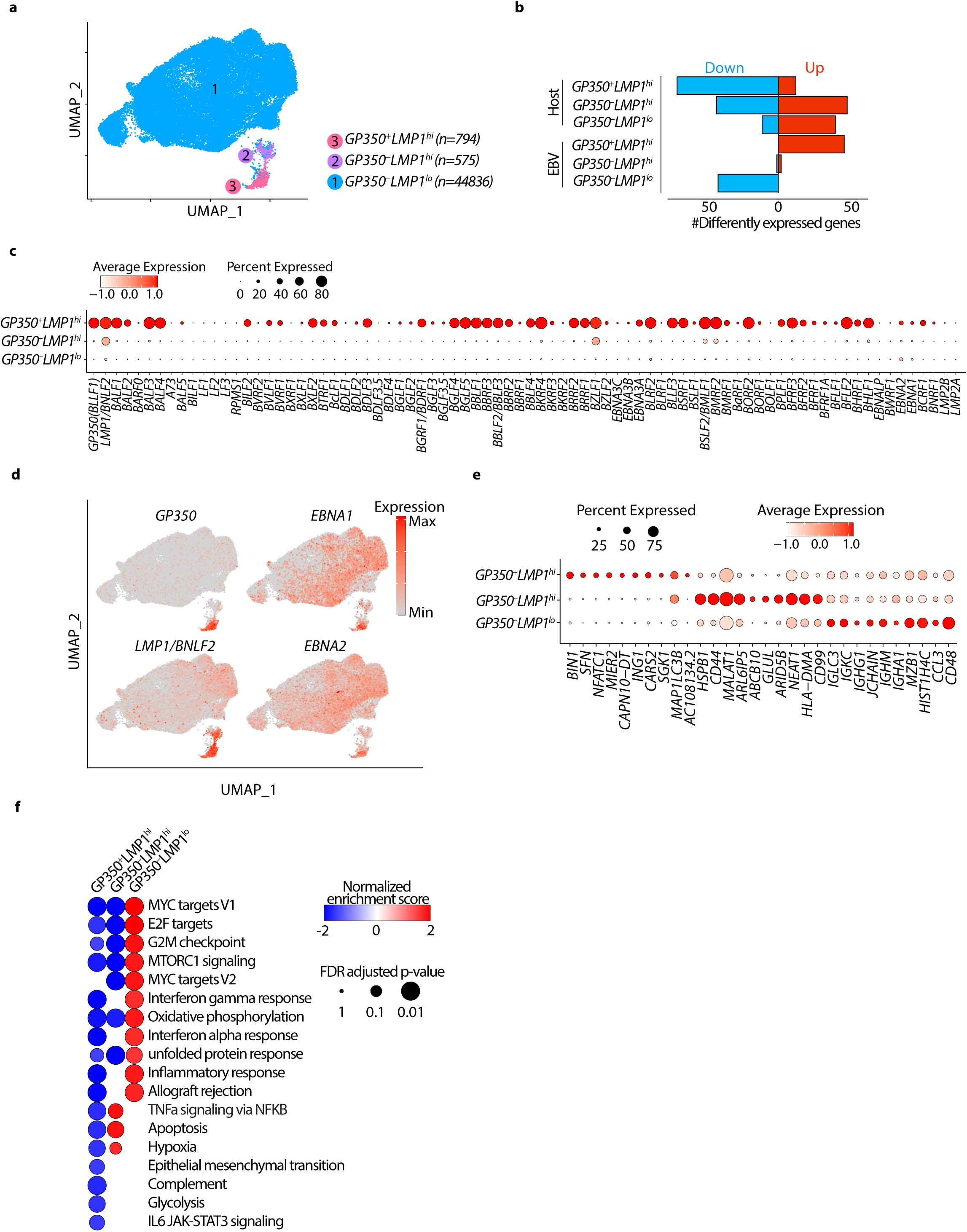
Single cell RNA-sequencing analyses resolve LCLs into three distinct populations. (**a**) Integrated UMAP showing 3 major cell types w.r.t. EBV status in nine LCLs used in this study. (**b**) Numbers of differentially expressed genes (FC>1.5 and adjusted p-val < 0.05) in indicated cluster compared to other clusters. (**c-d**) mRNA expression of EBV genes across all clusters shown as dot plot (**c**) or projected on the UMAP (**d**). (**e**) mRNA expression of top 10 human host cell defining genes across all clusters. (**f**) Significantly enriched hallmark pathways by GSEA comparing transcriptomes of cells in indicated cluster with all other cells. The positive and negative enrichment scores indicate activation and inactivation of the indicated pathway in each cell cluster, respectively. Only pathways that are enriched (FDR<5%) in at least one of the clusters are shown.

*GP350*^−^*LMP1*^*lo*^ cells comprised 94-98% of all LCLs across all the samples (**Figs. 1a, S1a-b**). They displayed minimal expression of *LMP1/BNLF2* and minimal or no expression of EBV lytic genes, including *GP350, BMRF1, BALF1* and *BALF3* (**Fig. 1c**). Additionally, these cells expressed latency genes, including *EBNA1* and *EBNA2*, indicating that this cluster mainly consisted of transformed cells that were in the EBV latent phase (**Fig. 1d**). This cluster was also the highest expressor of genes from immunoglobulin heavy or light chains, indicating the mature status of these transformed B cells (**Figs. 1e, S1g**). Consistently, nearly a quarter of cells in this cluster expressed high levels of *PRDM1*, indicating that these cells might have entered plasmacytic differentiation (41).

The two lytic clusters, *GP350*^−^*LMP1*^*hi*^ and *GP350*^+^*LMP1*^*hi*^ cells, respectively, each accounted for 1-5% of all LCLs (**Figs. 1a, S1b**). The salient viral feature of both clusters was the high expression of *LMP1/BNLF2*, a gene with well-established contribution to oncogenic human B-cell transformation (42) and *BZLF1. GP350*^+^*LMP1*^*hi*^ cells were the highest expressors of EBV lytic genes, including *GP350, BZLF1* and *BMRF1*, while *GP350*^−^*LMP1*^*hi*^ cells express very few lytic genes (**Fig. 1c**). Remarkably, these two clusters had distinct expressions of host genes (**Figs. 1e, S1f**). Consistent with previous reports (33, 34), *GP350*^*+*^*LMP1*^*hi*^ cells highly expressed host *NFATC1, MIER2, SFN* and *SGK1* genes and were the highest expressors of host box-dependent myc-interacting protein 1 (*BIN1*). Conversely, *GP350*^*–*^*LMP1*^*hi*^ cells had the highest expression of host genes *HSPB1, ABCB10, MALAT1* and *CD44* (**Fig. 1e**).

To obtain insights into the functional state of cells in each cluster, we performed geneset enrichment analysis (GSEA), comparing the transcriptomes of cells in each cluster against those of cells from the other two clusters (**Fig. 1f**), and querying enrichment of all 50 hallmark genesets curated by the Molecular Signatures Database (MSigDB) (43). Genes differently regulated in *GP350*^*–*^*LMP1*^*lo*^ cells were significantly enriched in MYC targets, MTORC1 signaling and inflammatory response. Conversely, genes differently regulated in *GP350*^*–*^*LMP1*^*hi*^ cells were enriched in tumor necrosis factor alpha signaling, apoptosis and hypoxia (**Fig. 1f**). As expected, genes differently regulated in *GP350*^*+*^*LMP1*^*hi*^ cells were significantly depleted of genesets from most hallmark pathways, including MYC targets, MTORC1 signaling and interferon responses (**Figs. 1f, S1h**). This is consistent with the fact that fully lytic EBV reactivation pauses transcription of most host genes and pathways, which is evidenced by significantly reduced numbers of total host transcripts in lytic cells (**Fig. S1i**).

Collectively these analyses indicated that LCLs are predominantly comprised of three distinct cell populations characterized by differences in expression of both host and viral genes, notably cells containing EBV in the latent phase (*GP350*^*–*^*LMP1*^*lo*^), cells containing virus in the lytic phase (*GP350*^*+*^*LMP1*^*hi*^) and cells that were in between lytic and latent phases (*GP350*^*–*^ *LMP1*^*hi*^). Furthermore, these data suggested that distinct functional states of individual LCL clusters may be related to expression of genes encoded by EBV and the host cell.

### GP350^−^LMP1^hi^ LCLs have a HIF1A-associated signature

We next explored the transcriptional regulators of host gene expression. Our attention was drawn to *HIF1A* because *GP350*^*–*^*LMP1*^*hi*^ cells were high expressors of several genes including *HSPB1, MALAT1* and *CD44* (**Fig. 1e**) that in other settings are known to be regulated by hypoxia or HIF1-α (44-46) and because our GSEA analysis had also indicated that the transcriptomes of these cells are highly enriched in the hypoxia gene set (**Fig. 1f**). HIF1-α is a critically important TF that is tightly regulated by oxygen tension and transactivates many genes essential for cellular responses and adaptation to hypoxia (47). To better characterize *GP350*^*–*^ *LMP1*^*hi*^ cells, we therefore quantified the mRNA expression of *HIF1A*, the gene that encodes HIF1-α, and its classical direct target *PDL1* (48) in all clusters. *HIF1A* and *PDL1* were both more highly expressed in *GP350*^*–*^*LMP1*^*hi*^ cells compared to others (**Fig. 2a**). This was specifically evident for *HIF1A* as its expression levels were significantly higher in *GP350*^*–*^ *LMP1*^*hi*^ cells (**Figs. 2a, S2a**). To determine whether the changes in *HIF1A* expression could have any functional consequence, we next assessed the expression of HIF1-α target genes. We sourced a public list (49) of HIF1-α–induced (n=110) and HIF1-α–repressed (n=77) genes from MSigDB (50) and assessed the expression of these two sets in all 3 identified LCL clusters (**Fig. 2b** and **Table S2**). *GP350*^*–*^*LMP1*^*hi*^ cells had the highest and lowest expressions among all clusters for HIF1-α–induced and HIF1-α–repressed genes, expressed as the module score (51), respectively (**Fig. 2b**). We confirmed these findings using two additional independent lists of HIF1-α–regulated genes (52) (**Fig. S2b-c** and **Table S2**).

**Figure 2.**
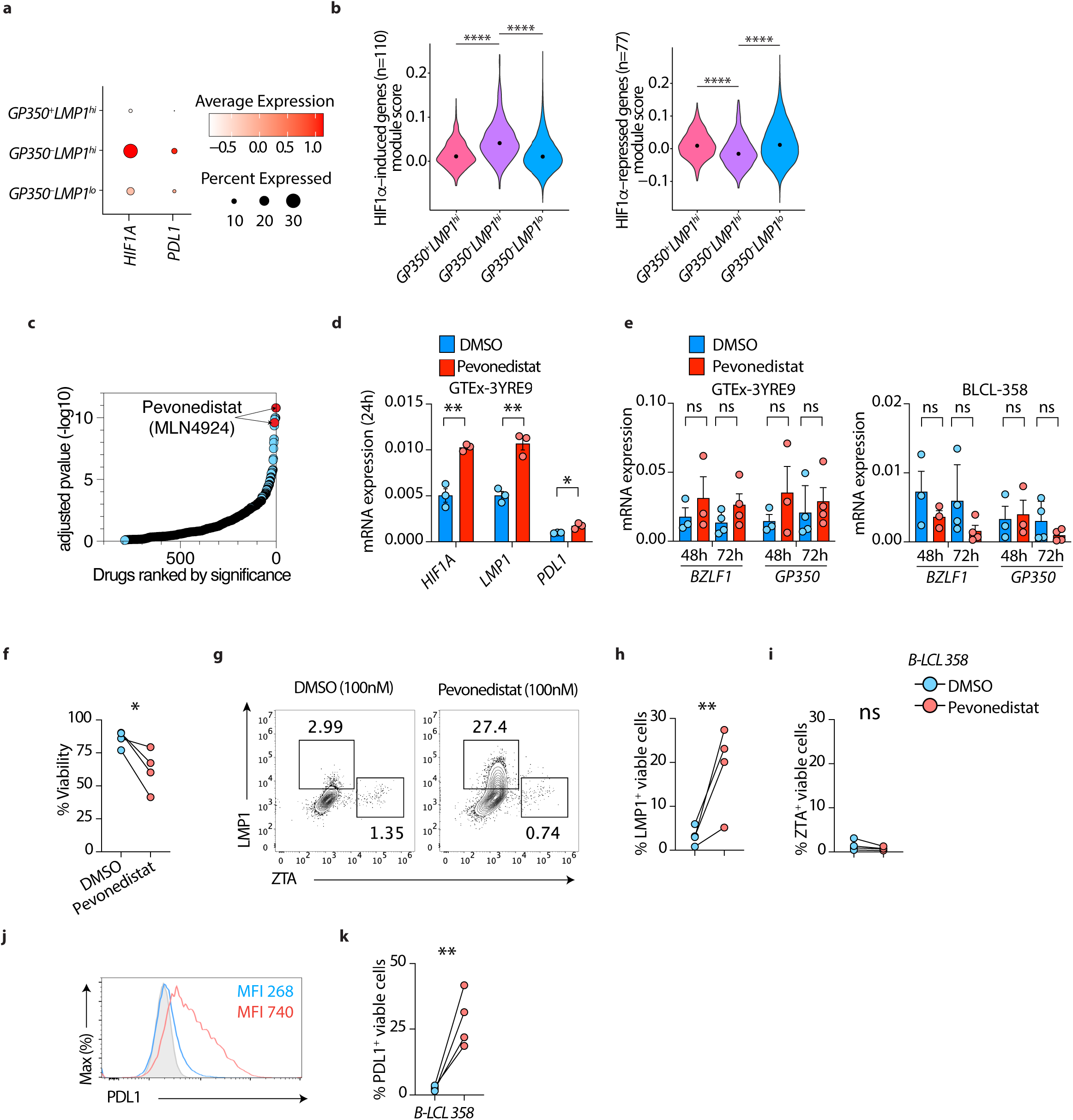
*GP350*^*–*^*LMP1*^*hi*^ LCLs have a HIF1A-associated signature. **(a)** mRNA expression of *HIF1A* or *CD274* genes across all clusters as dot plot. **(b)** Module score of HIF1A induced genes (left panel) or HIF1A-repressed genes (right panel). HIF1A induced and repressed genes are sourced from MSigDB (M1308). **** p<0.0001 by two-tailed Wilcoxon rank-sum test. **(c)** Enrichr based drugs predicted (out of 906 total drugs) to counteract genes induced in *GP350*^*–*^ *LMP1*^*hi*^ LCLs compared to other cells, ordered by adjusted p-value. Drugs are sourced from Enrichr library “Drug_Perturbations_from_GEO_down”. (**d**-**e)** mRNA expression of indicated host (**d**) or EBV (**e**) genes in LCLs treated with 100nM DMSO or Pevonedistat. UBC was used as a housekeeping gene. (**f-k**) flow cytometry on BLCL-358 treated with 100nM DMSO or Pevonedistat for 72hr. Plots showing cell viability and LMP1, BZLF1 or PDL1 expression in LCLs treated with DMSO or Pevonedistat. Shown are cumulative %viability plots (**f**), representative flow cytometry plots (**g**) and cumulative data showing %LMP1^+^ **(h)** and % BZLF1^+^(**i**) in gated live LCLs. (**j-k**) Representative PDL1 expression as mean fluorescent intensity (MFI) or cumulative %PDL1^+^ in gated live LCLs. Data in (**d-k**) are from n=3 or n = 4 independent experiments; gating strategy is shown in **Fig. S2d**. * p<0.05; ** p<0.01; *** p<0.001; **** p<0.0001 by two-tailed paired ratio t-test.

We next predicted pharmaceutical agents that could induce the unique gene signatures of cells in the *GP350*^*–*^*LMP1*^*hi*^ cells, using methods established previously by our group (32, 53). Among the topmost significant drugs predicted to enhance host gene expression pattern of *GP350*^*–*^*LMP1*^*hi*^ cells was Pevonedistat (MLN4924) (**Fig. 2c**). Pevonedistat is a ubiquitin-activating enzyme E1 inhibitor that significantly stabilizes HIF1-α to potentiate its function (54). Because the HIF1-α pathway was one of the main features of *GP350*^*–*^*LMP1*^*hi*^ cells, we hypothesized that enhancing HIF1-α signaling might preferentially induce this program. To test this hypothesis, we treated LCLs with Pevonedistat and measured *HIF1A, LMP1, PDL1* and GP350. HIF1-α potentiation markedly induced mRNA expression of *HIF1A, LMP1* and *PDL1* (**Fig. 2d**), but not *GP350* or *BZLF1* (**Fig. 2e**). To further substantiate these observations at the single cell level, and confirm expression of protein, we treated three different LCLs with Pevonedistat and performed flow cytometry. Pevonedistat reduced cell viability by nearly 30% (**Figs. 2f, S2e**). The frequency of LMP1^+^ cells was significantly increased (**Figs. 2g-h, S2f-g**) without increasing that of ZTA (**Figs. 2i, S2h)**. The frequency of PDL1^+^ cells and PDL1 expression were also significantly increased upon treatment (**Fig. 2j-k, S2i**). We have also performed dose titration of Pevonedistat and have observed dose dependent increase of LMP1 and PD-L1, but not BZLF1, in gated live cells (**Fig. S3**), suggesting that Pevonedistat preferentially induce LMP1^+^ cells without increasing full lytic cycle.

### GP350^−^LMP1^lo^ LCLs have distinct MYC-dependent transcriptional programs

We next focused on transcriptional regulators of *GP350*^*–*^*LMP1*^*lo*^ LCLs, the cluster containing EBV in the latent phase. MYC-regulated genes were among the top affected pathways when comparing transcriptomes of LCL clusters against each other (**Fig. 1f**) and box-dependent myc-interacting protein 1 (*BIN1*) was one of the top host genes distinguishing *GP350*^*+*^ from *GP350*^*–*^ cells (**Fig. 1e**). Moreover, MYC has been described as a key host factor repressing EBV lytic reactivation (55). Thus, we further examined the role of MYC in shaping the distinct LCL clusters. Because MYC is a transcription factor, we first determined the fraction of differently expressed genes in each cluster directly bound by MYC. For this, we sourced a publicly available ChIP-seq dataset for MYC in GM12878 (GSM822290, curated by ENCODE). Nearly 18-24% of genes differently expressed in each cluster were directly bound by MYC, with *GP350*^*+*^*LMP1*^hi^ cluster having the most number of MYC targets (**Fig. 3a** and **Table S2**). This was significantly higher than what would be expected by chance because only ∼10% of all human genes are bound by MYC in GM12878 (**Fig. Sa**).

**Figure 3.**
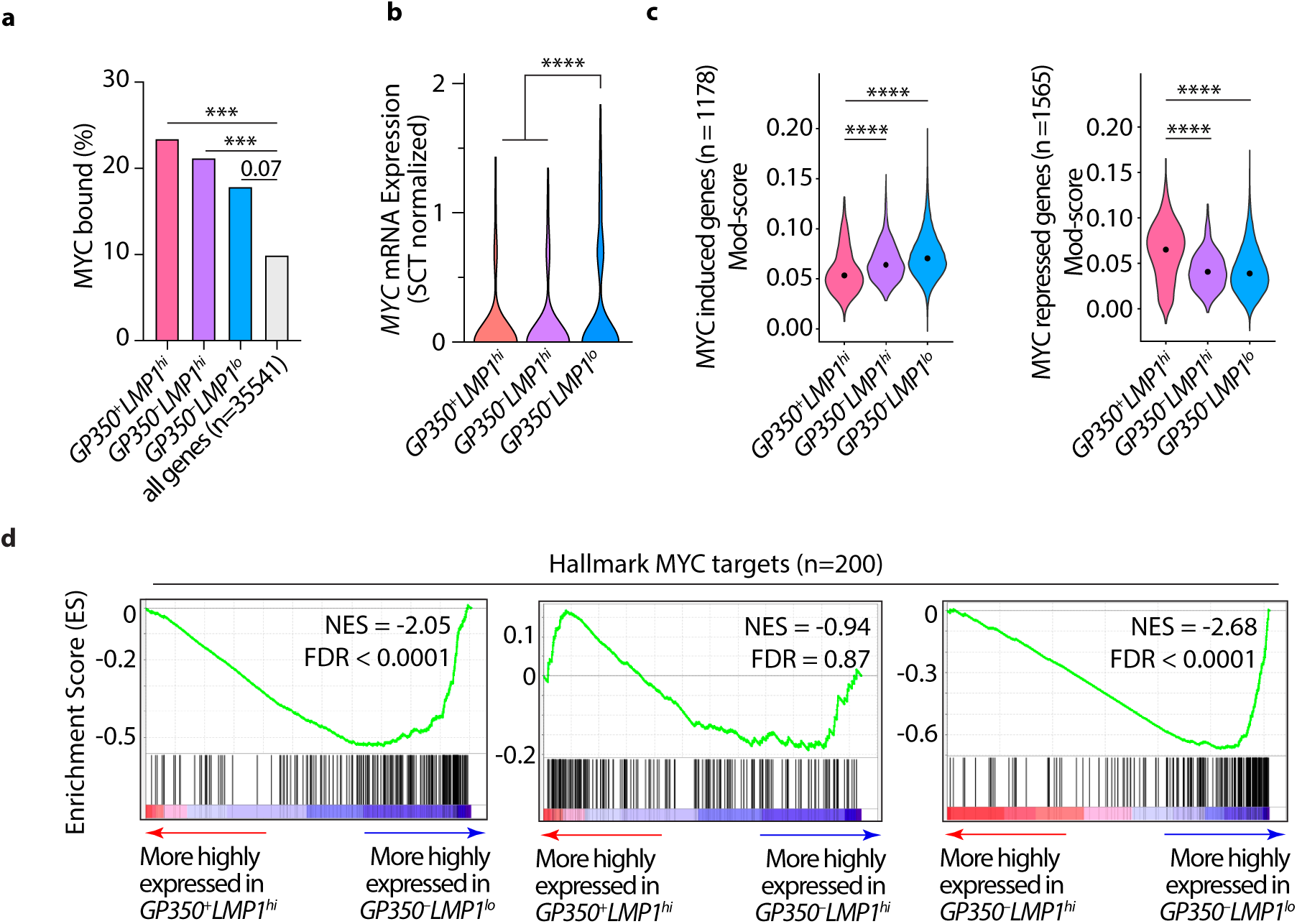
GP350^−^LMP1^lo^ LCLs have distinct MYC-dependent transcriptional regulation. **(a)** fraction of differentially expressed genes (**Fig. 1b**) from each indicated cluster or all human genes (n=35541) that are bound by MYC. MYC bound genes (n=3534) in GM12878 are obtained from (81). *** p<0.001 by fisher exact test. **(b)** MYC mRNA expression across all clusters. **** p<0.0001 by two-tailed Wilcoxon rank-sum test. **(c)** Module score of MYC induced genes (left panel) or MYC-repressed genes (right panel). **** p<0.0001 by two-tailed Wilcoxon rank-sum test. MYC induced or repressed genes are sourced from (55). **(d)** GSEA plots comparing transcriptomes of indicated clusters for the enrichment in hallmark of MYC targets. NES: Normalized enrichment scores. FDR: False Discovery Rate.

The mRNA expression of *MYC* was significantly higher in *GP350*^*–*^*LMP1*^*lo*^ LCLs than in either of the other two clusters (**Fig. 3b**). To determine whether MYC is biologically active, we looked for the signature of genes regulated by MYC. We curated a list of genes regulated by MYC in GM12878 from a publicly available dataset (55) (**Table S2**). Expression of MYC-induced genes was significantly higher (**Fig. 3c**, left panel) and MYC-repressed genes significantly lower (**Fig. 3c**, right panel) in *GP350*^*–*^*LMP1*^*lo*^ cells than in the other two clusters. We also performed GSEA comparing transcriptomes of cells from each cluster against MYC targets curated by MSigDB (43). Consistent with our earlier observation (**Fig. 1f**), genes that were more highly expressed in *GP350*^*–*^*LMP1*^*lo*^ cells were highly enriched in MYC targets (**Fig. 3d**, left and right panels), while there was no significant difference between LMP1^hi^ clusters (**Fig. 3d**, middle panel). Collectively, these data indicated that MYC preferentially regulates a subset of genes that are differently expressed in *GP350*^*–*^*LMP1*^*lo*^ LCLs.

### Super-enhancer-regulated genes are less highly expressed in GP350^+^LMP1^hi^ LCLs

EBV-infected cells periodically enter the lytic phase to produce progeny viruses but in EBV-immortalized lymphoblastoid cells EBV is mostly in the latent state. Earlier studies have shown that a small percentage of these cells are lytic (30). However, due to the technical challenges at the time, it was difficult to distinguish the cells in lytic phase from cells at latency phase in a mixed population. The recent development of scRNA-seq techniques allows us to capture the cells in lytic phase together with their transcriptome.

Nearly 10% of genes are regulated by multiple enhancers forming a complex architecture known as “super-enhancers” (SEs). SE-regulated genes are critically important for cell identity (27) and are associated with both Mendelian and polygenic diseases (56, 57) as well as cancers (58). Enhancer-promoter interactions are the cornerstones of mammalian gene regulation. We have previously shown that EBV episomes make reproducible contacts with the human genome at SE loci (23). To explore whether EBV disrupts modes of gene regulation in the three LCL subsets, we sourced a list of 257 annotated SE regulated genes from GM12878 (26) and determined whether these genes are differently expressed in the three identified LCL clusters. Unexpectedly, expression of SE-regulated genes, summarized as the module score, was significantly lower in *GP350*^*+*^*LMP1*^*hi*^ cells, the cluster containing EBV in the lytic state, than in the other two subsets (**Fig. 4a**). We observed similar results when we used an independent curated set of 187 EBV-associated SEs (22) (**Fig. S4a**). Examples of such genes included *MYC*, which contains one of the largest SEs in the genome (22), *IRF4, RUNX3, PAX5, IKZF3* and *DUSP22* (**Fig. 4b**). These findings suggested that lytic EBV is associated with disruption of expression of host genes regulated through SEs. To explore this possibility, we performed GSEA analysis comparing the transcriptomes of *GP350*^*+*^*LMP1*^*+*^ cells to cells from the other two clusters for enrichment of all SE-regulated genes. This orthogonal approach also indicated that genes less highly expressed in *GP350*^*+*^*LMP1*^*+*^ cells compared to the cells in the other two clusters were markedly enriched in SE-regulated genes (**Fig. S4b**).

**Figure 4.**
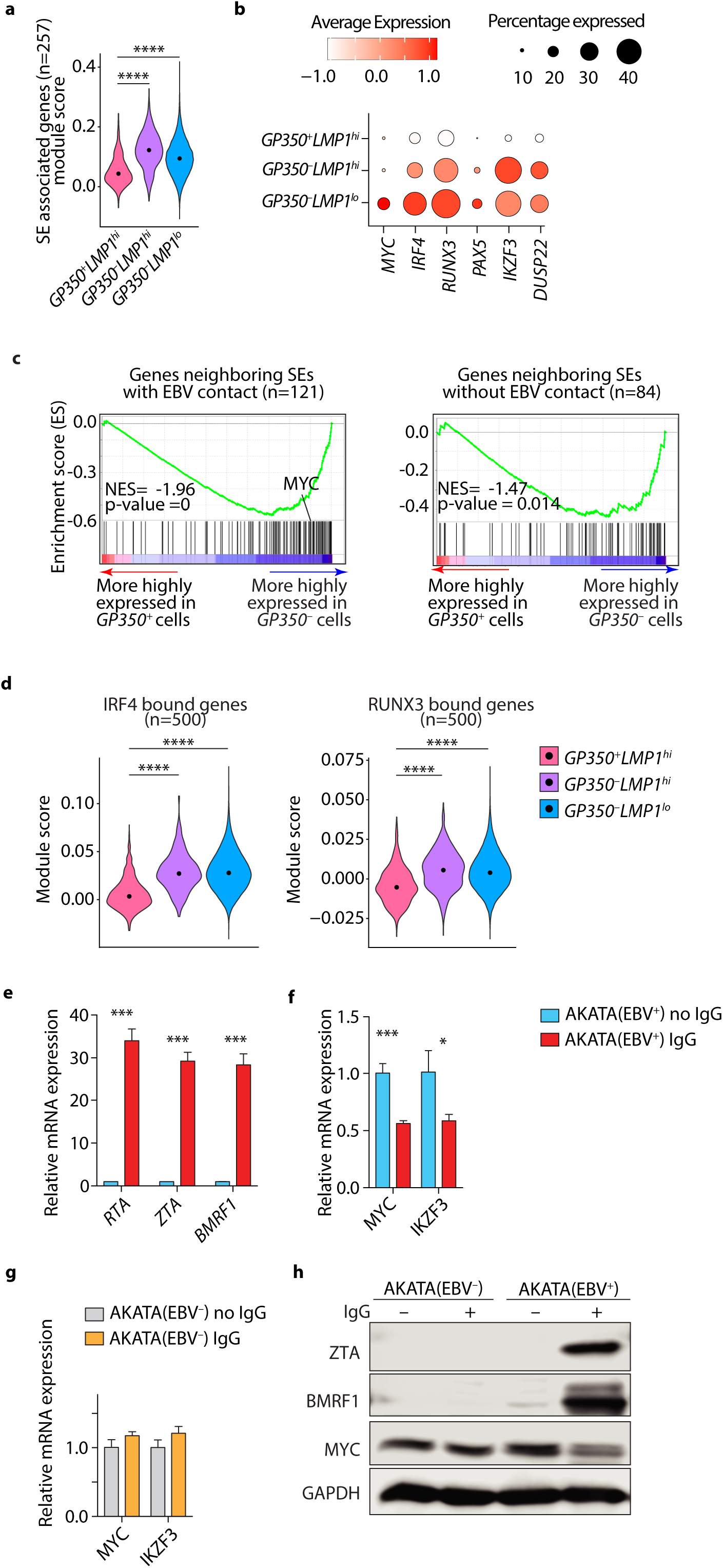
Super-enhancer-regulated genes are less highly expressed in *GP350*^*+*^ LCLs. **(a)** Module score of SE containing genes in GM12878. SEs and their annotation are sourced from (26). **(b)** mRNA expression of select SE containing genes across all cell types. **(c)** GSEA plots comparing transcriptomes of *GP350*^+^ and *GP350*^*–*^ cells for enrichment in genes neighboring SEs with (left panel) or without (right panel) EBV contacts. NES: Normalized enrichment scores. **(d)** Module score of n=500 top IRF4 (left panel) and RUNX3 (right panel) bound genes. **** p<0.0001 by two-tailed Wilcoxon rank-sum test. **(e-f)** mRNA expression of indicated EBV (**e**) or host (**f**) genes in EBV^+^-AKATA cells with or without anti-IgG (1:200) treatment after 48 hours. (**g**) Control EBV^−^-AKATA cells are included when measuring host genes. Data are from n=3 independent experiments. * p<0.05; ** p<0.01; *** p<0.001 by two-tailed unpaired t-test. **(h)** Western blots of lysates of AKATA cells treated with or without anti-IgG after 48 hours. Shown are representative (n=2) images of ZTA (BZLF1), BMRF1 and MYC with GAPDH as loading control.

We next assessed whether genes in these clusters were differently expressed when their associated SEs physically interact with EBV episomes. To this end, we divided SE-regulated genes into those that physically interact, or not, with EBV episomes and performed GSEA analysis comparing *GP350*^*+*^ cells to the cells of other two *GP350*^*–*^ subsets. Genes that were more highly expressed in *GP350*^*–*^ cells were significantly enriched in SEs that interact with EBV episomes (**Fig. 4c**, left panel). This enrichment was less evident for SEs that do not interact with EBV episomes (**Fig. 4c**, right panel). To determine the functional consequences of differential expression of SE-regulated genes across LCL clusters, we focused on the transactivator IRF4 and transcription factor RUNX3 for which we could source their direct targets from ChIP-seq experiments and assess the expression of their targets. We noted that expression of both *IRF4-* and *RUNX3*-bound genes, summarized as the module score, was significantly lower in *GP350*^*+*^*LMP1*^*hi*^ cells (**Figs. 4d, S4c**), in which expression of both these TFs was also the lowest (**Fig. 4b**).

Since *GP350*^*+*^*LMP1*^*hi*^ cells represented the cluster in which lytic reactivation of EBV was apparent (**Fig. 1e**), we tested whether EBV reactivation affects the expression of SE containing genes. To this end, we treated EBV^+^ AKATA cells with either anti-IgG or carrier. Anti-IgG is a potent inducer of EBV lytic reactivation in these cells (59). After stimulation, we measured mRNA and/or protein expression of EBV lytic markers and the host SE-regulated gene *MYC* (**Figs. 4e-h**). As expected, anti-IgG induced strong expression of the EBV lytic markers *RTA, ZTA* and *BMRF1* (**Fig. 4e**). In contrast, the expression of both *MYC* and *IKZF3* were significantly repressed following anti-IgG-treatment of cells (**Fig. 4f**). This effect was specifically a predicate of EBV-reactivation since anti-IgG did not repress *MYC* or *IKZF3* expression in EBV^−^ AKATA cells (**Fig. 4g**). These observations were further confirmed by immunoblots of ZTA, BMRF1 and MYC proteins (**Fig. 4h**). Consistently, a recent study has shown that depletion of MYC reactivates the EBV lytic cycle (55). To test whether depletion of IRF4 can similarly reactivate EBV lytic cycle, we reanalyzed RNA-seq from GM12878 LCLs that were subjected to IRF4 deletion via the CRISPR/Cas9 system (29). In this setting, depletion of *IRF4* induced multiple EBV lytic genes, including *GP350, RTA, ZTA* and *BMRF1* (**Fig. S4d**). Consistently, a recent study has found that IRF4 knockdown in LCLs induces EBV lytic reactivation in LCLs and lytically infected cells have increased NFATc1 and decreased IRF4 expression (34). Collectively, our data suggest that SE-regulated genes are less highly expressed in *GP350*^+^*LMP1*^*hi*^ LCLs, which show evidence of lytic EBV reactivation, and that experimental induction of EBV lytic cycle also represses expression of these genes.

### Disruption of super-enhancers in LCLs induces EBV lytic reactivation

The reciprocal relationship between expression of SE-regulated genes in *GP350*^*+*^*LMP1*^*hi*^ LCLs and EBV reactivation suggested the possibility that SEs may be necessary for maintenance of EBV in the latent phase. To test this possibility, we initially selected SEs near *MYC* and *IRF4/DUSP22* and performed CRISPR-mediated knockout or inactivation and then measured the expression of EBV lytic markers. For these experiments, appropriate guide RNAs were situated within the SEs at sites of maximal H3K27ac signal, a marker of active regions of the genome, especially promoters and enhancers. The selected sites were bound by one or more viral transcription factors (e.g. EBNA2, LP, 3A and 3C) and/or host NF-κB family members (e.g. RelA, RelB, cRel, p50 and p52) and interacted within a topologically associated domain that contained the SE (**Figs. 5a, 5c**). Dual sgRNAs targeting both sides of *MYC* SE (∼525 kb upstream) successfully deleted *MYC* SE from the genome (**Fig. S5a**), which led to a reduction of *MYC* transcription and upregulation of EBV lytic genes, namely ZTA, RTA, BGLF5 and BMRF1 expression (**Fig. 5b**). Similarly, inactivation of *IRF4* SE by CRISPR-dcas9 tethered with a repressor consisting of KRAB and the transcription repression domain of MeCP2 successfully reduced *IRF4* SE activity (**Fig. S5b**). This led to decrease in *IRF4/DUSP22* mRNA expression and a significant increase of EBV lytic gene expression two days post inactivation **(Fig. 5d)**. Deletion of both *MYC* and *IRF4* genes have been previously shown to induce EBV lytic phase. Unexpectedly, however, when we measured the expression of EBV lytic genes at earlier time points, we observed that EBV lytic genes were significantly induced prior to decrease of *IRF4* (**Fig. S5c**). This suggests the possibility that SE might be also necessary for the maintenance of EBV latency. To further explore this possibility, we selected another SE in the same topological associated domain of MIR155HG (**Fig. 5e**), using the same criteria as above and performed CRISPR-mediated inactivation. The disruption of this SE did not significantly reduce the expression of *MIR155HG*; however, it significantly increased the expression of EBV lytic genes (**Fig. 5f**), namely *ZTA, RTA, BGLF5* and *BMRF1*. CRISPRi disruption of RUNX3 SE also had similar activity (**Fig. 5g, h**). Collectively these data indicate that deletion of select host SEs leads to lytic reactivation of EBV and, by extension, that host SEs, or their target genes, are necessary for maintenance of EBV in the latent phase.

**Figure 5.**
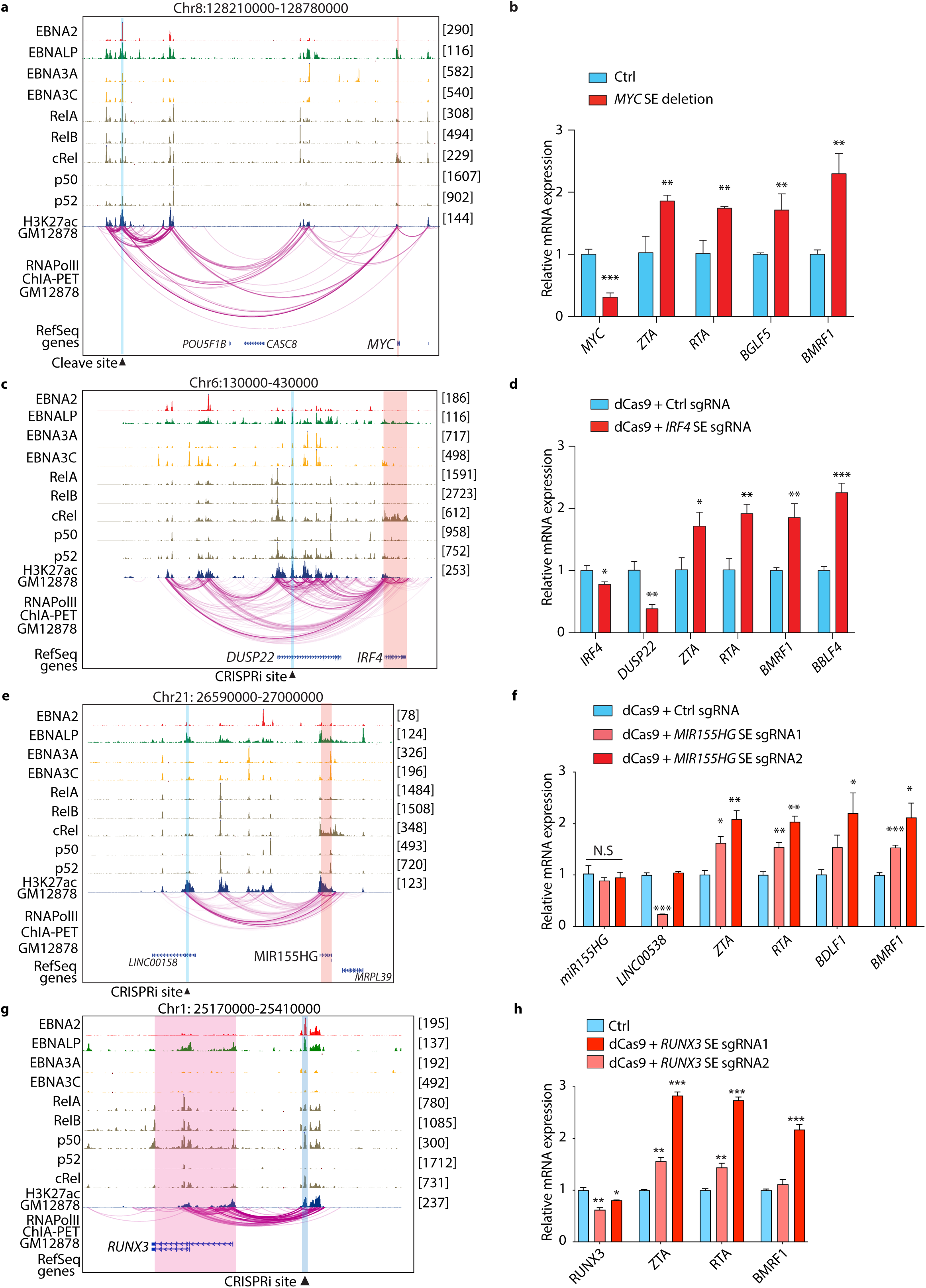
Disruption of super-enhancers in LCLs induces EBV lytic reactivation. (**a**,**c**,**e**,**g)** Genome browser tracks showing EBV transcription factors EBNA2, EBNALP, EBNA3A and EBNA 3C, host transcription factors RELA, RELB, c-REL, p50 and p52 and H3K27ac at the *MYC* **(a)**, *IRF4/DUSP22* **(c)**, *MIR155HG* **(e)** and *RUNX3* (**g**) loci. The CRISPR cleavage site or inactivation site is highlighted with vertical blue box. The expected affected target genes are highlighted by vertical red box. **(b**,**d**,**f**,**h)** mRNA expression of select host and EBV genes after CRISPR mediated knockout (**b**) or inactivation (**d**,**f**,**h**) of indicated site (bottom black triangle in **a**,**c, e** or **g**) in GM12878 cells. Data are from n=3 independent experiments. * p<0.05, ** p<0.01, *** p<0.001 by two-tailed unpaired t-test.

## Discussion

LCLs have been instrumental for genetic and functional studies of human diseases over the past several decades (60). We and others have previously analyzed large numbers of LCL bulk RNA-seq data and found that EBV lytic gene expression correlates with cellular cancer-associated pathways, such as interferon-alpha, WNT and B cell receptor signaling (4, 30). However, these data were generated from bulk populations of cells, which biases insights towards those occurring in the largest sub-populations. While the majority of LCLs contain EBV in the latent phase of its life cycle, a small fraction (<5%) demonstrate spontaneous EBV reactivation, indicating that LCLs as a whole are heterogeneous. Important aspects of LCL heterogeneity have recently been explored using single cell RNA-sequencing (33). This analysis focused on heterogeneity within and across LCLs with respect to immunoglobulin isotypes, which further associated with pathways involving activation and differentiation of B cells. We have taken an integrative approach to combine the same data with several more datasets that are generated across different conditions and eliminate batch and technical effects. This integrative analysis provides a consistent representation of the data for downstream analyses and thus has the potential to uncover previously undetected biology. Specifically, we found LCLs to have higher heterogeneity in relation to the EBV status than previously appreciated. Specifically, we identified three prominent clusters that were marked by the expression of the EBV genes *GP350* and *LMP1/BNLF2*. Of note, the EBV genome has extensive numbers of overlapping genes such as *LMP1* and *BNLF2a/b*, making the quantification of their mRNA expression more challenging (61). This challenge could be further exacerbated by the 3’ mRNA capture bias in some of the current single cell technologies. Nevertheless, we showed that these clusters have distinct transcriptional programs and identified MYC and HIF1-α as transcriptional regulators of gene expression. LCLs in the *GP350*^+^ cluster expressed SE-regulated genes at significantly lower levels compared to cells in the other two clusters. Physical interactions between SE-containing loci and EBV episomes marked genes in *GP350*^+^ LCLs that were particularly lowly expressed. Indeed, in proof-of-principle experiments we found that experimental lytic reactivation of EBV disrupted expression of SE-regulated genes and, conversely, that disruption of SEs induced EBV lytic reactivation. For *IRF4* or *MIR155HG* associated SEs, lytic reactivation after SE disruption occurred prior to *IRF4* downregulation, suggesting that these SEs themselves might be necessary for the maintenance of EBV latency. However, further studies are needed to fully discern this observation.

In the largest subset of LCLs, annotated as *GP350*^*–*^*LMP1*^*lo*^, EBV was clearly in the latent phase. This cluster showed a host gene signature enriched in MYC-regulated targets. As an oncogene, *MYC* is exquisitely carefully regulated by an archetypal SE. MYC itself binds to the EBV genome origin of lytic replication and suppresses DNA looping to the promoter of the lytic cycle initiator gene *BZLF1* (55). MYC depletion reactivates the lytic cycle in different cells (55). Consistent with this, when we deleted the *MYC* SE, MYC expression was decreased and EBV lytic genes were induced, supporting the role of MYC as a repressor of EBV lytic activation. Thus, it appears that both the *MYC* gene and its associated SE have a role in maintenance of EBV latency.

Another GP350 negative cluster, characterized by high expression of *LMP1/BNLF2*, was the highest expressor of several host genes including *HSPB1, MALAT1* and *CD44* that are known features of cancer stem cells (62-64) and escape from apoptosis (65, 66). Interestingly, LMP1 alone induces CSC features in nasopharyngeal cell lines (67). However, such characteristics have not previously been ascribed to LCLs and warrant further investigation. LMP1 is a known oncogene and expressed in most EBV associated cancers (68) and has been previously associated with synthesis of HIF1-α protein and its DNA binding activity (69). Here, we also found that *GP350*^*–*^*LMP1*^*hi*^ LCLs have a prominent HIF1-α signature and could be preferentially induced by Pevonedistat. These cells also expressed higher frequencies of PDL1, which were markedly increased upon Pevonedistat treatment. Interestingly, a recent study has found an association between numbers of PDL1 expressing B cells and the development of AIDS related non-Hodgkin lymphoma (70). Such PDL1 expressing B cells have previously been described to suppress effector function of immune cells (71). Thus, the identification of these cells might play an important role in understanding the oncogenesis and may suggest that drugs that stabilize HIF1-α might inadvertently induce LMP1 in diseases associated with EBV type III latency programs such as AIDS-associated B cell lymphoma, post-transplant lymphoproliferative disorder and diffuse large B cell lymphoma.

B cell differentiation into plasma cell has been linked to EBV lytic replication (72, 73). Specifically, PRDM1, a known driver of B cell differentiation into plasma cells (74), promotes EBV lytic replication by activating the transcription from immediate early gene promoters of ZTA and RTA (75). A recent single cell RNA-seq analysis of LCLs have revealed a positive correlation between specific immunoglobulin isotype and cell differentiation markers (33). However, these immunoglobin genes were not specifically characterized in lytic cells. We have found that the mRNA expression of *PRDM1* and a range of immunoglobulin genes in *GP350*^*+*^*LMP1*^hi^ cells was lower than in latent cells, which contrasts with previous reports about the role of *PRDM1* in EBV lytic reactivation (**Fig. S1e**). It is possible that *PRDM1* is important for initiation, but not maintenance, of the lytic cycle. Another possibility is that the transcription factor activity, but not the overall expression level, of PRDM1 is important for lytic replication. Further study is clearly required to delineate this relationship.

In summary, we performed integrative analysis of publicly available single cell RNA-seq data from different LCLs to help resolve their heterogeneity. We identified a novel cluster of cells that are between lytic and latent stage, marked by LMP1 and controlled by HIF1a. We also found that the mRNA expression of super-enhancer target genes is inversely correlated with lytic status of the cells and consistently CRISPR perturbation of super-enhancers increased the expression of EBV lytic genes. Our studies revealed EBV associated heterogeneity among LCLs that contribute to EBV life cycle and biology.

## Acknowledgements

This work was supported by extramural research programs of the NIH (R35GM138283 to MK and 5R01AI123420 and 5R01CA047006 to BZ) and the Showalter Trust (research award to MK). This research was supported (in part) by the Intramural Research Programs of the National Institute of Diabetes and Digestive and Kidney Diseases (project number ZIA/DK075149 to BA). The authors also gratefully acknowledge the SIRG Graduate Research Assistantships Award to BY and SC and support from the Purdue University Center for Cancer Research, P30CA023168.

## Author contributions

BY, ZZ, SK, LW and MJ performed computational work. CW, SC, IS, YZ, YH, SS, KS and NA performed experimental work. All other authors contributed significantly to computational, experimental and/or conceptual development of this work. BZ and MK conceptualized the study, supervised the project and wrote the manuscript.

## Competing interests

The authors have no competing interests to declare.

## Methods and Materials

### Cell culture

LCL-358 (catalog no. 1038-3754NV17, Astarte Biologics), GTEX-UPJH-0001-SM-3YRE9, GM12878, AKATA EBV positive and AKATA EBV negative cells were cultured in RPMI1640 (VWRL0105-0500) media supplemented with 10% fetal calf serum (Gibco or Hyclone), 100 unit/mL streptomycin and 100 mg/mL penicillin (Gibco or Life Technologies). HEK293T cells purchased from ATCC were cultured in Dulbecco modified Eagle medium supplemented with 10% fetal calf serum (Gibco), 100 unit/mL streptomycin and 100 mg/mL penicillin. All the cells were maintained at 37 °C in a 5% CO_2_ humidified chamber. Cells were routinely confirmed to be mycoplasma negative according to PCR Mycoplasma Detection Kit (ABM Inc.) and were used at low passage (<10) number but were not independently authenticated.

### CRISPRi repression

Plasmid dCAS9-KRAB-MeCP2 (#110821) purchased from Addgene was packaged with lentiviruses and used to transduce LCLs for 2 days followed by selection with 5ug/ml blasticidin for another 5 days. The expression of dCAS9-KRAB-MeCP2 was verified by western blot. sgRNAs targeting genomic sites of interest were designed with online software Benchling (www.benchling.com). sgRNAs were annealed and cloned into LentiGuide-Puro vector according to previously published protocol (76). LentiGuide-Puro containing sgRNAs were packaged into lentiviruses and were used to transduce LCLs stably expressing dCAS9-KRAB-MeCP2. Cells were selected with 3ug/ml puromycin for 3 days and then allowed to grow for another 2 days. The list of sgRNAs are provided in **Table S3**

### qRT-PCR

Cells were harvested and washed once with cold PBS. Total mRNAs were extracted using PureLink RNA mini kit (Life Technologies) or Direct-zol RNA extraction kit with DNase I treatment (Zymo Research) following manufacturer’s instructions. mRNAs were then reverse transcript into cDNA with iScript™ Reverse Transcription Supermix (Bio-rad) or OneScript Plus cDNA Synthesis SuperMix (ABM Inc.). cDNAs were amplified on an CFX96 Touch real-time PCR detection system (Bio-Rad) and SYBG Green (Thermo Fisher) was used to detect cDNA amplification. All experiments were performed in triplicates in total reaction volumes of 15 μL using BrightGreen 2X qPCR MasterMix-No Dye (ABM Inc.). A housekeeping gene was used to normalize gene expression. RNA relative expression was calculated using the 2 ΔΔCT method. The value for the cells transduced with non-targeting sgRNA was set to 1. The list of all qPCR probes is provided in **Table S3**.

### ChIP-qPCR

LCLs stably expressing dCAS9-KRAB-MeCP2 were transduced with lentiviruses expressing sgRNAs. Two days after transduction cells were selected with 3ug/ml puromycin for another 3 days. Cells were then collected and fixed with 1% formaldehyde. The cells were lysed and sonicated with bioruptor (Diagenode). Sonicated chromatin was diluted and precleared with protein A beads followed by incubation with 4ug H3K27ac (Abcam, #ab4729) or control antibodies with rotating at 4°C overnight. The next day, Protein A/salmon DNA beads (Millipore, #16-157) were used to capture protein–DNA complexes. After precipitation, beads were washed with low salt wash buffer (1% TritonX-100, 0.1% SDS, 2mM EDTA (pH8.0), 150mM NaCl, 20mM Tris-HCl (pH 8.0)) once, high salt wash buffer (1% TritonX-100, 0.1% SDS, 2mM EDTA (pH8.0), 500mM NaCl, 20mM Tris-HCl (pH 8.0)) twice, Licl wash buffer (0.25M LiCl, 1% NP-40, 1% NaDOC, 1mM EDTA, 10mM Tris-HCl (pH 8.0)) once, and TE buffer (1mM EDTA, 10mM Tris-HCl (pH 8.0)) once. Each wash was performed by gently spinning down beads at 300g for one minute and re-suspend beads with 1ml wash buffer followed by shaking at 4°C for 5 minutes. DNA and protein complexes were eluted with elution buffer (1% SDS, 100mM NaHCO3). Protein–DNA complexes were reverse cross-linked with proteinase K (Thermo Fisher, #EO0491). DNA was purified by using QIAquick Spin columns (Qiagen, #28104). qPCR was used to quantify the DNA from ChIP assay and normalize it to the percent of input DNA.

### Induction of EBV

AKATA EBV positive and negative cells were treated with IgG (Agilent, # A042301-2) at a final concentration of 0.5% followed by incubation at 37°C, 5% CO2 for 6 hrs. Cells were then centrifuged and re-suspended with fresh RPMI1640 supplemented with 10% FBS and continue culture for another 48 hrs. mRNAs were extracted by using PureLink RNA mini kit (Life Technologies), qRT-PCR was used to detect EBV lytic gene expression. To induce LMP1 expression, LCLs were treated with 100 nM of NEDD8 inhibitor -MLN4924 (Pevonedistat) (A gift from Dr. Liu) or DMSO control and were collected at indicated time points for qRT-PCR and/or Flow cytometry.

### Dual CRISPR mediated DNA deletion

Dual gRNAs were designed with webtools from benchling (www.benchling.com) and were cloned into pLentiGuide-Puro (Addgene Plasmid #52963) using the Multiplex gRNA kit (System Biosciences) according to the manufacturer protocol. The success of gRNAs insertion was verified by sequencing with U6 promoter primer. HEK293T cells were used to package lentiviruses by co-transfecting viral packaging plasmids pCMV-VSV-G (Addgene #8454), psPAX2 (Addgene #12260) and the pLentiGuide-Puro vector containing the target sgRNAs. 18 hrs after transfection, media were changed to fresh RPMI media containing 30% of FBS. 24 and 48hrs later, supernatant containing lentivirus was collected and filtered with a 0.45 micron filter. LCLs in which Cas9 was stably expressed were transduced with the filtered lentivirus (Day 0) for 2 days, and then selected with 3ug/mL Puromycin for 3 days. On Day 5, genomic DNA was extracted using the DNeasy Blood & Tissue kit (Qiagen) and RNA was extracted with the PureLink RNA Mini kit (Ambion). Genomic deletions were verified by PCR using the PrimeSTAR polymerase (Clonetech). qRT-PCR was done using the Power SYBR Green RNA-to-CT 1-Step Kit (Applied Biosystems).

### Single-cell RNA sequencing analysis

10x Genomics Cell Ranger 6.0.2 count (77) was used to align the raw sequencing reads to a customized human (GRCh38) and EBV (NC_007605, obtained from (61)) hybrid reference genome to generate barcode and UMI counts. Seurat (v4) (78) was applied for the downstream analysis and visualization of the data as following: Genes that were expressed in less than 3 cells were discarded. Cells with >20% of their unique molecular identifiers (UMIs) mapping to mitochondrial genes or cells with <250 detected genes were discarded. Only cells with >80% log10 (Genes per UMI) were retained. Cell cycle score for each cell was calculated by Seurat function CellCycleScorin using human cell cycle genes. SCTransform was then used to normalize the dataset using default parameters while regressing out mitochondrial genes and cell cycle scores (S and G2M) and identify variable genes. Doublets were removed by R package DoubletFinder (v2.0) (79). Then, the datasets were integrated based on “anchors” identified among datasets (nfeatures = 2000, normalization.method = “SCT”) prior to perform linear dimensional reduction by Principal Component Analysis (PCA). The top 50 PCs were included in a Uniform Manifold Approximation and Projection (UMAP) dimensionality reduction. Clusters were identified on a shared nearest neighbor (SNN) graph the top 50 PCs with the Louvain algorithm using three resolutions (i.e., 0.1, 0.3, 0.5). the clusters corresponding to latent EBV life cycle were combined as one cluster for the downstream analyses. Differential gene expression was determined by “FindMarkers” function on SCT normalized expression values with the default Wilcox Rank Sum test either as one versus rest or as a direct comparison with default parameters except logfc.threshold = 0. The cell annotation was based on the EBV genes and top differentially expressed genes. Gene list module scores were calculated with Seurat function AddModuleScore (51). This calculates the average scaled expression levels of each gene list, subtracted by the expression of control feature sets (n= 100). All the displayed expression values on violin plots, feature plots and dot plots are SCT normalized expression values. The IRF4 bound genes are sourced from the ChIP-Atlas “Target Genes” database (80) with options: “hg38” as the genome and “+/-5kb” as distance from TSS. Target genes with binding score not less than 500 in GM12878 cells are selected. All genesets used in this study are provided in **Table S2**.

### Gene set enrichment analysis (GSEA)

GSEA was performed using pre-ranked mode and “No Collapse” options. The pre-ranked gene lists were ranked by the SCT normalized expression fold-change between comparison groups. EBV-contacted and EBV-non-contacted genesets are curated from our previous study (23) and provided in **Table S2**.

### Statistical analysis and data visualization

Statistical analyses were performed using GraphPad PRISM 9 (La Jolla, CA, USA) with the method detailed in the legend.

### Flow cytometry

All stained/fixed samples were acquired on Attune NxT Flow Cytometer (Thermo Fisher Scientific) with necessary internal controls to help assign gates. Fluorescence from multiple antibodies were compensated using AbC Total Compensation beads (Thermo Fisher Scientific, catalog no. A10497). In all experiments, cells were collected and stained with fixable viability dye eFluor780 (1:2000 dilution Life Technologies, catalog no. 65-0865-14;) followed by surface staining for PDL1 (CD274; clone 29E.2A3; BioLegend catalog no. 329714; 1:60 dilution) as per the manufacturer’s instructions. Cells were then fixed with 4% methanol-free formaldehyde (Thermo Fisher Scientific, catalog no. 28908) followed by intracellular staining for BZLF1 (Santa Cruz Biotechnology, catalog no. sc-53904; 1:60 dilution) and LMP1 (clone LMPO24; Novus Biologicals, catalog no. NBP2-50383; 1:60 dilution) using FoxP3/Transcription factor staining buffer set (eBioscience, catalog no. 5523) as per manufacturer’s instructions. Data were analyzed using FlowJo and cumulated using GraphPad PRISM software.

## Data sources and availability

The single cell RNA-seq data are sourced from GSE126321 for GM12878 and GM18502; GSE111912 for GM12891; GSE158275 for LCL777B958, LCL777M81 and LCL461B958; GSE162528 for LCL; and GSE121926 for GM22648 and GM22649. The ChIA-PET in GM12878 is from GSE127053. The ChIP-seq data are from EBNA2: GSE29498; EBNALP: GSE49338; EBNA3A: GSM1429820; EBNA3C: GSE52632; NF-κB: GSE55105 and H3K27Ac: GSM733771.

## References

1. Epstein MA, Achong BG, Barr YM. 1964. Virus Particles in Cultured Lymphoblasts from Burkitt’s Lymphoma. Lancet 1:702–3.

2. Cohen JI, Fauci AS, Varmus H, Nabel GJ. 2011. Epstein-Barr virus: an important vaccine target for cancer prevention. Sci Transl Med 3:107fs7.

3. Howley PM, Knipe DM, Cohen JL, Damania BA. 2021. Fields Virology: DNA Viruses. Wolters Kluwer Health.

4. Chakravorty S, Yan B, Wang C, Wang L, Quaid JT, Lin CF, Briggs SD, Majumder J, Canaria DA, Chauss D, Chopra G, Olson MR, Zhao B, Afzali B, Kazemian M. 2019. Integrated Pan-Cancer Map of EBV-Associated Neoplasms Reveals Functional Host-Virus Interactions. Cancer Res 79:6010–6023.

5. Cameron JE, Fewell C, Yin Q, McBride J, Wang X, Lin Z, Flemington EK. 2008. Epstein-Barr virus growth/latency III program alters cellular microRNA expression. Virology 382:257–66.

6. Young LS, Yap LF, Murray PG. 2016. Epstein-Barr virus: more than 50 years old and still providing surprises. Nat Rev Cancer 16:789–802.

7. Lieberman PM. 2014. Virology. Epstein-Barr virus turns 50. Science 343:1323–5.

8. Pope JH, Horne MK, Scott W. 1968. Transformation of foetal human keukocytes in vitro by filtrates of a human leukaemic cell line containing herpes-like virus. Int J Cancer 3:857–66.

9. Knipe DM, Howley PM. 2013. Fields virology, 6th ed. Wolters Kluwer/Lippincott Williams & Wilkins Health, Philadelphia, PA.

10. Pei Y, Wong JH, Robertson ES. 2020. Herpesvirus Epigenetic Reprogramming and Oncogenesis. Annu Rev Virol 7:309–331.

11. Frappier L. 2015. Ebna1. Curr Top Microbiol Immunol 391:3–34.

12. Saridakis V, Sheng Y, Sarkari F, Holowaty MN, Shire K, Nguyen T, Zhang RG, Liao J, Lee W, Edwards AM, Arrowsmith CH, Frappier L. 2005. Structure of the p53 binding domain of HAUSP/USP7 bound to Epstein-Barr nuclear antigen 1 implications for EBV-mediated immortalization. Mol Cell 18:25–36.

13. Wu H, Ceccarelli DF, Frappier L. 2000. The DNA segregation mechanism of Epstein-Barr virus nuclear antigen 1. EMBO Rep 1:140–4.

14. Kaiser C, Laux G, Eick D, Jochner N, Bornkamm GW, Kempkes B. 1999. The protooncogene c-myc is a direct target gene of Epstein-Barr virus nuclear antigen 2. J Virol 73:4481–4.

15. Portal D, Zhao B, Calderwood MA, Sommermann T, Johannsen E, Kieff E. 2011. EBV nuclear antigen EBNALP dismisses transcription repressors NCoR and RBPJ from enhancers and EBNA2 increases NCoR-deficient RBPJ DNA binding. Proc Natl Acad Sci U S A 108:7808–13.

16. Maruo S, Zhao B, Johannsen E, Kieff E, Zou J, Takada K. 2011. Epstein-Barr virus nuclear antigens 3C and 3A maintain lymphoblastoid cell growth by repressing p16INK4A and p14ARF expression. Proc Natl Acad Sci U S A 108:1919–24.

17. Skalska L, White RE, Parker GA, Turro E, Sinclair AJ, Paschos K, Allday MJ. 2013. Induction of p16(INK4a) is the major barrier to proliferation when Epstein-Barr virus (EBV) transforms primary B cells into lymphoblastoid cell lines. PLoS Pathog 9:e1003187.

18. Paschos K, Smith P, Anderton E, Middeldorp JM, White RE, Allday MJ. 2009. Epstein-barr virus latency in B cells leads to epigenetic repression and CpG methylation of the tumour suppressor gene Bim. PLoS Pathog 5:e1000492.

19. Laherty CD, Hu HM, Opipari AW, Wang F, Dixit VM. 1992. The Epstein-Barr virus LMP1 gene product induces A20 zinc finger protein expression by activating nuclear factor kappa B. J Biol Chem 267:24157–60.

20. Okabe A, Huang KK, Matsusaka K, Fukuyo M, Xing M, Ong X, Hoshii T, Usui G, Seki M, Mano Y, Rahmutulla B, Kanda T, Suzuki T, Rha SY, Ushiku T, Fukayama M, Tan P, Kaneda A. 2020. Cross-species chromatin interactions drive transcriptional rewiring in Epstein-Barr virus-positive gastric adenocarcinoma. Nat Genet 52:919–930.

21. McClellan MJ, Wood CD, Ojeniyi O, Cooper TJ, Kanhere A, Arvey A, Webb HM, Palermo RD, Harth-Hertle ML, Kempkes B, Jenner RG, West MJ. 2013. Modulation of enhancer looping and differential gene targeting by Epstein-Barr virus transcription factors directs cellular reprogramming. PLoS Pathog 9:e1003636.

22. Zhou H, Schmidt SC, Jiang S, Willox B, Bernhardt K, Liang J, Johannsen EC, Kharchenko P, Gewurz BE, Kieff E, Zhao B. 2015. Epstein-Barr virus oncoprotein super-enhancers control B cell growth. Cell Host Microbe 17:205–16.

23. Wang L, Laing J, Yan B, Zhou H, Ke L, Wang C, Narita Y, Zhang Z, Olson MR, Afzali B, Zhao B, Kazemian M. 2020. Epstein-Barr Virus Episome Physically Interacts with Active Regions of the Host Genome in Lymphoblastoid Cells. J Virol 94.

24. Buschle A, Mrozek-Gorska P, Cernilogar FM, Ettinger A, Pich D, Krebs S, Mocanu B, Blum H, Schotta G, Straub T, Hammerschmidt W. 2021. Epstein-Barr virus inactivates the transcriptome and disrupts the chromatin architecture of its host cell in the first phase of lytic reactivation. Nucleic Acids Res 49:3217–3241.

25. Tempera I, Klichinsky M, Lieberman PM. 2011. EBV latency types adopt alternative chromatin conformations. PLoS Pathog 7:e1002180.

26. Hnisz D, Abraham BJ, Lee TI, Lau A, Saint-Andre V, Sigova AA, Hoke HA, Young RA. 2013. Super-enhancers in the control of cell identity and disease. Cell 155:934–47.

27. Whyte WA, Orlando DA, Hnisz D, Abraham BJ, Lin CY, Kagey MH, Rahl PB, Lee TI, Young RA. 2013. Master transcription factors and mediator establish super-enhancers at key cell identity genes. Cell 153:307–19.

28. Jiang S, Zhou H, Liang J, Gerdt C, Wang C, Ke L, Schmidt SCS, Narita Y, Ma Y, Wang S, Colson T, Gewurz B, Li G, Kieff E, Zhao B. 2017. The Epstein-Barr Virus Regulome in Lymphoblastoid Cells. Cell Host Microbe 22:561–573 e4.

29. Ma Y, Walsh MJ, Bernhardt K, Ashbaugh CW, Trudeau SJ, Ashbaugh IY, Jiang S, Jiang C, Zhao B, Root DE, Doench JG, Gewurz BE. 2017. CRISPR/Cas9 Screens Reveal Epstein-Barr Virus-Transformed B Cell Host Dependency Factors. Cell Host Microbe 21:580–591 e7.

30. Arvey A, Tempera I, Tsai K, Chen HS, Tikhmyanova N, Klichinsky M, Leslie C, Lieberman PM. 2012. An atlas of the Epstein-Barr virus transcriptome and epigenome reveals host-virus regulatory interactions. Cell Host Microbe 12:233–45.

31. Yan B, Chakravorty S, Mirabelli C, Wang L, Trujillo-Ochoa JL, Chauss D, Kumar D, Lionakis MS, Olson MR, Wobus CE, Afzali B, Kazemian M. 2021. Host-Virus Chimeric Events in SARS-CoV-2-Infected Cells Are Infrequent and Artifactual. J Virol 95:e0029421.

32. Yan B, Freiwald T, Chauss D, Wang L, West E, Mirabelli C, Zhang CJ, Nichols EM, Malik N, Gregory R, Bantscheff M, Ghidelli-Disse S, Kolev M, Frum T, Spence JR, Sexton JZ, Alysandratos KD, Kotton DN, Pittaluga S, Bibby J, Niyonzima N, Olson MR, Kordasti S, Portilla D, Wobus CE, Laurence A, Lionakis MS, Kemper C, Afzali B, Kazemian M. 2021. SARS-CoV-2 drives JAK1/2-dependent local complement hyperactivation. Sci Immunol 6.

33. SoRelle ED, Dai J, Bonglack EN, Heckenberg EM, Zhou JY, Giamberardino SN, Bailey JA, Gregory SG, Chan C, Luftig MA. 2021. Single-cell RNA-seq reveals transcriptomic heterogeneity mediated by host-pathogen dynamics in lymphoblastoid cell lines. Elife 10.

34. Bristol JA, Brand J, Ohashi M, Eichelberg MR, Casco A, Nelson SE, Hayes M, Romero-Masters JC, Baiu DC, Gumperz JE, Johannsen EC, Dinh HQ, Kenney SC. 2022. Reduced IRF4 expression promotes lytic phenotype in Type 2 EBV-infected B cells. PLoS Pathog 18:e1010453.

35. Osorio D, Yu X, Yu P, Serpedin E, Cai JJ. 2019. Single-cell RNA sequencing of a European and an African lymphoblastoid cell line. Sci Data 6:112.

36. Ozgyin L, Horvath A, Hevessy Z, Balint BL. 2019. Extensive epigenetic and transcriptomic variability between genetically identical human B-lymphoblastoid cells with implications in pharmacogenomics research. Sci Rep 9:4889.

37. Sokka J, Yoshihara M, Kvist J, Laiho L, Warren A, Stadelmann C, Jouhilahti EM, Kilpinen H, Balboa D, Katayama S, Kyttala A, Kere J, Otonkoski T, Weltner J, Trokovic R. 2022. CRISPR activation enables high-fidelity reprogramming into human pluripotent stem cells. Stem Cell Reports 17:413–426.

38. Zhang X, Li T, Liu F, Chen Y, Yao J, Li Z, Huang Y, Wang J. 2019. Comparative Analysis of Droplet-Based Ultra-High-Throughput Single-Cell RNA-Seq Systems. Mol Cell 73:130–142 e5.

39. Osorio D, Yu X, Zhong Y, Li G, Yu P, Serpedin E, Huang JZ, Cai JJ. 2019. Single-Cell Expression Variability Implies Cell Function. Cells 9.

40. Hafemeister C, Satija R. 2019. Normalization and variance stabilization of single-cell RNA-seq data using regularized negative binomial regression. Genome Biol 20:296.

41. Tunyaplin C, Shaffer AL, Angelin-Duclos CD, Yu X, Staudt LM, Calame KL. 2004. Direct repression of prdm1 by Bcl-6 inhibits plasmacytic differentiation. J Immunol 173:1158–65.

42. Kaye KM, Izumi KM, Kieff E. 1993. Epstein-Barr virus latent membrane protein 1 is essential for B-lymphocyte growth transformation. Proc Natl Acad Sci U S A 90:9150–4.

43. Liberzon A, Birger C, Thorvaldsdottir H, Ghandi M, Mesirov JP, Tamayo P. 2015. The Molecular Signatures Database (MSigDB) hallmark gene set collection. Cell Syst 1:417–425.

44. Salle-Lefort S, Miard S, Nolin MA, Boivin L, Pare ME, Debigare R, Picard F. 2016. Hypoxia upregulates Malat1 expression through a CaMKK/AMPK/HIF-1alpha axis. Int J Oncol 49:1731–6.

45. Liang G, Li S, Du W, Ke Q, Cai J, Yang J. 2017. Hypoxia regulates CD44 expression via hypoxia-inducible factor-1alpha in human gastric cancer cells. Oncol Lett 13:967–972.

46. Whitlock NA, Agarwal N, Ma JX, Crosson CE. 2005. Hsp27 upregulation by HIF-1 signaling offers protection against retinal ischemia in rats. Invest Ophthalmol Vis Sci 46:1092–8.

47. Wang GL, Jiang BH, Rue EA, Semenza GL. 1995. Hypoxia-inducible factor 1 is a basic-helix-loop-helix-PAS heterodimer regulated by cellular O2 tension. Proc Natl Acad Sci U S A 92:5510–4.

48. Noman MZ, Desantis G, Janji B, Hasmim M, Karray S, Dessen P, Bronte V, Chouaib S. 2014. PD-L1 is a novel direct target of HIF-1alpha, and its blockade under hypoxia enhanced MDSC-mediated T cell activation. J Exp Med 211:781–90.

49. Gross C, Dubois-Pot H, Wasylyk B. 2008. The ternary complex factor Net/Elk-3 participates in the transcriptional response to hypoxia and regulates HIF-1 alpha. Oncogene 27:1333–41.

50. Subramanian A, Tamayo P, Mootha VK, Mukherjee S, Ebert BL, Gillette MA, Paulovich A, Pomeroy SL, Golub TR, Lander ES, Mesirov JP. 2005. Gene set enrichment analysis: a knowledge-based approach for interpreting genome-wide expression profiles. Proc Natl Acad Sci U S A 102:15545–50.

51. Tirosh I, Izar B, Prakadan SM, Wadsworth MH, 2nd, Treacy D, Trombetta JJ, Rotem A, Rodman C, Lian C, Murphy G, Fallahi-Sichani M, Dutton-Regester K, Lin JR, Cohen O, Shah P, Lu D, Genshaft AS, Hughes TK, Ziegler CG, Kazer SW, Gaillard A, Kolb KE, Villani AC, Johannessen CM, Andreev AY, Van Allen EM, Bertagnolli M, Sorger PK, Sullivan RJ, Flaherty KT, Frederick DT, Jane-Valbuena J, Yoon CH, Rozenblatt-Rosen O, Shalek AK, Regev A, Garraway LA. 2016. Dissecting the multicellular ecosystem of metastatic melanoma by single-cell RNA-seq. Science 352:189–96.

52. Elvidge GP, Glenny L, Appelhoff RJ, Ratcliffe PJ, Ragoussis J, Gleadle JM. 2006. Concordant regulation of gene expression by hypoxia and 2-oxoglutarate-dependent dioxygenase inhibition: the role of HIF-1alpha, HIF-2alpha, and other pathways. J Biol Chem 281:15215–26.

53. McGregor R, Chauss D, Freiwald T, Yan B, Wang L, Nova-Lamperti E, Zhang Z, Teague H, West EE, Bibby J, Kelly A, Malik A, Freeman AF, Schwartz D, Portilla D, John S, Lavender P, Lionakis MS, Mehta NN, Kemper C, Cooper N, Lombardi G, Laurence A, Kazemian M, Afzali B. 2020. An autocrine Vitamin D-driven Th1 shutdown program can be exploited for COVID-19. bioRxiv doi:10.1101/2020.07.18.210161.

54. Ryu JH, Li SH, Park HS, Park JW, Lee B, Chun YS. 2011. Hypoxia-inducible factor alpha subunit stabilization by NEDD8 conjugation is reactive oxygen species-dependent. J Biol Chem 286:6963–70.

55. Guo R, Jiang C, Zhang Y, Govande A, Trudeau SJ, Chen F, Fry CJ, Puri R, Wolinsky E, Schineller M, Frost TC, Gebre M, Zhao B, Giulino-Roth L, Doench JG, Teng M, Gewurz BE. 2020. MYC Controls the Epstein-Barr Virus Lytic Switch. Mol Cell 78:653–669 e8.

56. Afzali B, Gronholm J, Vandrovcova J, O’Brien C, Sun HW, Vanderleyden I, Davis FP, Khoder A, Zhang Y, Hegazy AN, Villarino AV, Palmer IW, Kaufman J, Watts NR, Kazemian M, Kamenyeva O, Keith J, Sayed A, Kasperaviciute D, Mueller M, Hughes JD, Fuss IJ, Sadiyah MF, Montgomery-Recht K, McElwee J, Restifo NP, Strober W, Linterman MA, Wingfield PT, Uhlig HH, Roychoudhuri R, Aitman TJ, Kelleher P, Lenardo MJ, O’Shea JJ, Cooper N, Laurence ADJ. 2017. BACH2 immunodeficiency illustrates an association between super-enhancers and haploinsufficiency. Nat Immunol 18:813–823.

57. Vahedi G, Kanno Y, Furumoto Y, Jiang K, Parker SC, Erdos MR, Davis SR, Roychoudhuri R, Restifo NP, Gadina M, Tang Z, Ruan Y, Collins FS, Sartorelli V, O’Shea JJ. 2015. Super-enhancers delineate disease-associated regulatory nodes in T cells. Nature 520:558–62.

58. Loven J, Hoke HA, Lin CY, Lau A, Orlando DA, Vakoc CR, Bradner JE, Lee TI, Young RA. 2013. Selective inhibition of tumor oncogenes by disruption of super-enhancers. Cell 153:320–34.

59. Yuan J, Cahir-McFarland E, Zhao B, Kieff E. 2006. Virus and cell RNAs expressed during Epstein-Barr virus replication. J Virol 80:2548–65.

60. Hussain T, Mulherkar R. 2012. Lymphoblastoid Cell lines: a Continuous in Vitro Source of Cells to Study Carcinogen Sensitivity and DNA Repair. Int J Mol Cell Med 1:75–87.

61. Casco A, Gupta A, Hayes M, Djavadian R, Ohashi M, Johannsen E. 2022. Accurate Quantification of Overlapping Herpesvirus Transcripts from RNA Sequencing Data. J Virol 96:e0163521.

62. Tang D, Yang Z, Long F, Luo L, Yang B, Zhu R, Sang X, Cao G, Wang K. 2019. Long noncoding RNA MALAT1 mediates stem cell-like properties in human colorectal cancer cells by regulating miR-20b-5p/Oct4 axis. J Cell Physiol 234:20816–20828.

63. Fan GC. 2012. Role of heat shock proteins in stem cell behavior. Prog Mol Biol Transl Sci 111:305–22.

64. Xiong J, Li Y, Tan X, Fu L. 2020. Small Heat Shock Proteins in Cancers: Functions and Therapeutic Potential for Cancer Therapy. Int J Mol Sci 21.

65. Wang QM, Lian GY, Song Y, Huang YF, Gong Y. 2019. LncRNA MALAT1 promotes tumorigenesis and immune escape of diffuse large B cell lymphoma by sponging miR-195. Life Sci 231:116335.

66. Yasuda M, Tanaka Y, Fujii K, Yasumoto K. 2001. CD44 stimulation down-regulates Fas expression and Fas-mediated apoptosis of lung cancer cells. Int Immunol 13:1309–19.

67. Kondo S, Wakisaka N, Muramatsu M, Zen Y, Endo K, Murono S, Sugimoto H, Yamaoka S, Pagano JS, Yoshizaki T. 2011. Epstein-Barr virus latent membrane protein 1 induces cancer stem/progenitor-like cells in nasopharyngeal epithelial cell lines. J Virol 85:11255–64.

68. Ersing I, Bernhardt K, Gewurz BE. 2013. NF-kappaB and IRF7 pathway activation by Epstein-Barr virus Latent Membrane Protein 1. Viruses 5:1587–606.

69. Wakisaka N, Kondo S, Yoshizaki T, Murono S, Furukawa M, Pagano JS. 2004. Epstein-Barr virus latent membrane protein 1 induces synthesis of hypoxia-inducible factor 1 alpha. Mol Cell Biol 24:5223–34.

70. Epeldegui M, Conti DV, Guo Y, Cozen W, Penichet ML, Martinez-Maza O. 2019. Elevated numbers of PD-L1 expressing B cells are associated with the development of AIDS-NHL. Sci Rep 9:9371.

71. Khan AR, Hams E, Floudas A, Sparwasser T, Weaver CT, Fallon PG. 2015. PD-L1hi B cells are critical regulators of humoral immunity. Nat Commun 6:5997.

72. Crawford DH, Ando I. 1986. EB virus induction is associated with B-cell maturation. Immunology 59:405–9.

73. Laichalk LL, Thorley-Lawson DA. 2005. Terminal differentiation into plasma cells initiates the replicative cycle of Epstein-Barr virus in vivo. J Virol 79:1296–307.

74. Shaffer AL, Lin KI, Kuo TC, Yu X, Hurt EM, Rosenwald A, Giltnane JM, Yang L, Zhao H, Calame K, Staudt LM. 2002. Blimp-1 orchestrates plasma cell differentiation by extinguishing the mature B cell gene expression program. Immunity 17:51–62.

75. Reusch JA, Nawandar DM, Wright KL, Kenney SC, Mertz JE. 2015. Cellular differentiation regulator BLIMP1 induces Epstein-Barr virus lytic reactivation in epithelial and B cells by activating transcription from both the R and Z promoters. J Virol 89:1731–43.

76. Sanjana NE, Shalem O, Zhang F. 2014. Improved vectors and genome-wide libraries for CRISPR screening. Nat Methods 11:783–784.

77. Zheng GX, Terry JM, Belgrader P, Ryvkin P, Bent ZW, Wilson R, Ziraldo SB, Wheeler TD, McDermott GP, Zhu J, Gregory MT, Shuga J, Montesclaros L, Underwood JG, Masquelier DA, Nishimura SY, Schnall-Levin M, Wyatt PW, Hindson CM, Bharadwaj R, Wong A, Ness KD, Beppu LW, Deeg HJ, McFarland C, Loeb KR, Valente WJ, Ericson NG, Stevens EA, Radich JP, Mikkelsen TS, Hindson BJ, Bielas JH. 2017. Massively parallel digital transcriptional profiling of single cells. Nat Commun 8:14049.

78. Hao Y, Hao S, Andersen-Nissen E, Mauck WM, 3rd, Zheng S, Butler A, Lee MJ, Wilk AJ, Darby C, Zager M, Hoffman P, Stoeckius M, Papalexi E, Mimitou EP, Jain J, Srivastava A, Stuart T, Fleming LM, Yeung B, Rogers AJ, McElrath JM, Blish CA, Gottardo R, Smibert P, Satija R. 2021. Integrated analysis of multimodal single-cell data. Cell 184:3573–3587 e29.

79. McGinnis CS, Murrow LM, Gartner ZJ. 2019. DoubletFinder: Doublet Detection in Single-Cell RNA Sequencing Data Using Artificial Nearest Neighbors. Cell Syst 8:329–337 e4.

80. Oki S, Ohta T, Shioi G, Hatanaka H, Ogasawara O, Okuda Y, Kawaji H, Nakaki R, Sese J, Meno C. 2018. ChIP-Atlas: a data-mining suite powered by full integration of public ChIP-seq data. EMBO Rep 19.

81. Lee BK, Bhinge AA, Battenhouse A, McDaniell RM, Liu Z, Song L, Ni Y, Birney E, Lieb JD, Furey TS, Crawford GE, Iyer VR. 2012. Cell-type specific and combinatorial usage of diverse transcription factors revealed by genome-wide binding studies in multiple human cells. Genome Res 22:9–24.

